# The Participation and Motivations of Grant Peer Reviewers: A Comprehensive Survey

**DOI:** 10.1101/479816

**Authors:** Stephen A Gallo, Lisa A Thompson, Karen B Schmaling, Scott R Glisson

**Affiliations:** American Institute of Biological Sciences; Washington State University

**Keywords:** Peer Review, Participation, Sustainability, Research Funding, Grant Applications, Motivation, Survey

## Abstract

Scientific peer reviewers play an integral role in the grant selection process, yet very little has been reported on the levels of participation or the motivations of scientists to take part in peer review. AIBS developed a comprehensive peer review survey that examined the motivations and levels of participation of grant reviewers. The survey was disseminated to 13,091 scientists in AIBS’s proprietary database. Of the 874 respondents, 76% indicated they had reviewed grant applications in the last 3 years; however, the number of reviews was unevenly distributed across this sample. Higher review loads were associated with respondents who had submitted more grant proposals over this time period, some of whom were likely to be study section members for large funding agencies. The most prevalent reason to participate in a review was to give back to the scientific community (especially among frequent grant submitters) and the most common reason to decline an invitation to review was lack of time. Interestingly, few suggested that expectation from the funding agency was a motivation to review. Most felt that review participation positively influenced their careers through improving grantsmanship and exposure to new scientific ideas. Of those who reviewed, respondents reported dedicating 2-5% of their total annual work time to grant review and, based on their self-reported maximum review loads, it is estimated they are participating at 56%-89% of their capacity, which may have important implications regarding the sustainability of the system. Overall, it is clear that participation in peer review is uneven and in some cases near capacity, and more needs to be done to create new motivations and incentives to increase the future pool of reviewers.

## Introduction

Peer review is the primary selection mechanism for funding research projects, utilizing panels of subject matter experts to evaluate and rank research funding applications. However, this process has recently been characterized as being under stress, with increasing demands for reviewers’ time and expertise as well as increased calls for an unbiased, transparent, and high integrity process despite limited budgets for peer review (Gropp, 2017). While there has been a steady increase in data-driven literature on grant peer review in recent years, there are still many key operational parameters that have yet to be characterized. Particularly important to the sustainability of the peer review system, but a relatively neglected area of research, is discerning the levels of participation and the motivations of the reviewers in this process, across the scientific community. Relatively little research has focused on the voluntary scientific workforce conducting these reviews, yet developing a better description of these important stakeholders is crucial to gauging the current strains on the peer review system. In order to alleviate these stresses and improve the quality of the outcomes, policy makers must understand reviewer motivations and their limits to ensure that the peer review system can be sustained in the 21^st^ century.

At the most basic level, an account of review participation levels across scientists is needed. While recent reports have highlighted the extraordinary amount of time reviewers spend evaluating journal article submissions (Stahel and Moore, 2014; Arns 2014, Kovanis, 2016), no such data exist for grant reviewers. Although the rosters of many standing panels are known to include more senior scientists, it is not known how or if review participation (including non-standing panel reviews) is dependent on gender, age, career stage or even clinical versus basic research backgrounds. Basic descriptions of reviewers are important not only to understand levels of diversity in this population, but also to predict who is likely to engage with the review system, as reviewer recruitment has been reported to be a frequent problem for funding organizations (Schroter 2010). More research is needed to estimate the annual commitment of grant reviewers, similar to surveys that have estimated the average journal review burden in terms of number of papers reviewed and hours spent (Ware, 2008). Without this knowledge, it is unclear how one can know anything about the sustainability of the grant peer review system. Along these same lines, distributional disparities of this effort across the scientific community are also needed, similar to those reported for journal review that suggest great disparities in review load (Kovanis, 2016). As many funding agencies expect successful applicants to participate in several years of study section membership (Amero, 2015), it is very likely some reviewer populations are more burdened than others; nevertheless, there is not currently any data examining levels of participation across the larger community or across populations with recent grant success (or recent grant submission). Therefore, one of the first steps toward establishing the sustainability of peer review is to get a sense of the distribution of current workloads.

In addition to information on current levels of participation, it is also important to know whether current review workloads are sustainable. Some surveys of the National Institutes of Health (NIH) study section reviewers have shown that majority prefer a workload in which grant peer review takes up less than 5% of their annual worktime, with the rest preferring 5-10% (Rockey, 2015). To date, there are not any studies directly examining reviewer-indicated preferences compared to actual workloads, which would estimate the extent of review system stress. These types of estimates will likely vary considerably across reviewer populations if the workloads are as uneven as they have been shown to be for journal reviewers (Kovanis, 2016).

Finally, a better understanding of the motivations behind reviewer involvement is paramount to ensuring a healthy and viable future peer review system. Some have suggested that improved grantsmanship and exposure to new ideas may be significant potential motivators, while others have warned against material incentives for reviewers (Irwin, 2013, Squazzoni, 2013). However, there has only been one study on grant reviewer motivations, which surveyed reviewers funded from European agencies (Schroter, 2010). While this study suggested that a sense of service is a main motivator, no analogous explorations have been reported for grant review in the US. Moreover, the Schroter study did not examine whether the amount of review participation was influenced by reviewer’s motivations.

To explore this area, the American Institute of Biological Sciences (AIBS) developed a comprehensive peer review survey addressing these areas. AIBS is a national scientific organization that promotes the use of science to inform decision-making that advances biology for the benefit of science and society. AIBS has provided independent peer review services for a variety of research funding organizations and institutes for over 50 years, and has developed a proprietary database of scientists that was used to distribute the survey. AIBS has published an analysis of some of the results from this survey previously (Gallo 2018), although this publication focused only on the application of the review criteria and the perceptions of both reviewers and applicants. Through the administration of the survey, AIBS also gathered data documenting levels of reviewer participation and associated time commitments, factors that predict this participation as well as reviewer motivations to accept and decline invitations to review, with the hope that the resulting analysis will contribute to more well-informed rationale for guiding, informing, and refining future peer review practices.

## Survey Methodology

As described in a previous manuscript (Gallo 2018), a comprehensive questionnaire containing 60 questions was developed for the peer review survey (the full survey is available in the **Appendix B**). The survey was divided into 5 sections, three of which will be analyzed in this manuscript: (1) Demographics; (2) Grant Submission and Peer Review Experience and (3) Reviewer Attitudes toward Grant Review. The sections (a) Investigator Attitudes Toward Grant Review and (b) Peer Review Panel Meeting Proceedings were not analyzed in this study. Only questions relevant to reviewer motivation, participation, and experience were analyzed. The questions’ response choices were dichotomous (yes/no), multiple choice, or interval rating scale (1-5). For each question, respondents were given the option to select “no answer/prefer not to answer.” In addition, text boxes were provided at the end of each section to allow respondents to elaborate on their responses. Based on beta testing, it was estimated that the survey would take around 15 minutes to complete. The survey was reviewed by the Washington State University Office of Research Assurances (Assurance# FWA00002946) and granted an exemption from IRB review (IRB#15268; 45 CFR 46.101(b)(2)).

The survey was sent in September 2016 to approximately 13,091 individuals in our database who had either participated on a peer review panel that had been convened by the institute (36%) or served as a PI/investigator/collaborator/consultant on an application that had been reviewed through the institute (71%) or both (12%). Again, results from a portion of this survey’s questions (focusing on perceptions of criteria usage by applicants and reviewers) were published in a separate manuscript (Gallo 2018), while this manuscript deals with questions pertaining to reviewer motivations and participation levels, and therefore analyzed results from different questions. As described previously (Gallo 2018), the survey was administered through Limesurvey, which allowed for identifying responses to be prevented from being linked back to participants. Each invited respondent was assigned a token by the system to ensure they only completed the survey once, but the linkage between their identity and the token was blinded from our staff. The survey was open for 2 months and once closed, the responses were exported and analyzed using basic statistical packages. To be included in the analysis, respondents had to fully submit their survey (hitting submit button at the end) and provide an answer to question 2.e, 2.f, and 2.g, which were focused on whether the respondent participated in reviews, if so, how many and what type. Percentages for levels of non-response to a given question are provided.

Multiple regression analyses required transforming demographic information into dichotomous variables. Gender was coded as female=1, male=0. As there were multiple categories for race, but as Caucasian was the most often indicated, race was coded as non-Caucasian=1, Caucasian=0. Again, there were multiple categories for degree, but as PhDs were the most frequent, degree was coded as non-PhD=1, PhD=0. As most respondents’ organizational affiliations were indicated to be academia, organization was coded as non-academia=1, academia=0. Career stage information was collected as 4 categories, so it was coded as early and mid-career stages=1, late and emeritus career stages=0. Age, work week hours, number of grant submissions and number of journal reviews were represented by continuous numeric values. In general, open-ended maximal values (e.g. 7 or more) were replaced with the smallest maximal value (e.g. 7) and age ranges (e.g. 30-39) were replaced by the lowest value in the range (e.g. 30) for ease of use in the regression model. The type of variable is indicated in the tables below (e.g. categorical or continuous numeric variables) to avoid confusion. How categorical variables were coded are indicated in parentheses (e.g. Female=1).

Respondents estimated their total number of reviews per year, the mode and number of days of review, number of assignments, hours per assignment, and total hours per week (as well as their preferred maximums for these values). The total proportion of yearly hours dedicated to grant review (annual working hours based on 50 weeks/year) was calculated by adding three components (pre-meeting hours, meeting hours, and travel time) and multiplying by the number of reviews reported. Pre-meeting review hours were determined from respondents’ estimates of the average number of hours spent pre-meeting per assignment multiplied by the estimate of average number of assignments. Meeting hours were determined by the reported average number of days per panel meeting multiplied by a presumed 8 hour work day. Travel time was estimated at 8 hours for face-to-face meetings (4 hours each way); no travel time was added for those who reported virtual or internet reviews. As all of these values were required to make this estimate (and many indicated they had not reviewed at all recently); 342 respondents with missing data from this analysis had to be removed, for a total N of 529. Anonymized survey data are available at https://doi.org/10.6084/m9.figshare.8132453.v1.

## Results

### Survey Response Rate

As mentioned in our previous publication, 13,091 individuals received an invitation to participate in the survey, had a valid email address and were not administrative officials (Gallo 2018). Of those individuals, 1,231 responded (9.4% response rate). Of these, 874 fully submitted their responses and provided an answer to questions 2efg, which were focused on whether the respondent participated in reviews, if so, how many and what type. Thus, the rate of completed responses to these sections of the survey (2efg) was 6.7%. All percentages are based on the total 874 respondents (unless otherwise noted).

### Survey Respondent Demographics

Overall, most respondents were Caucasian males over 40 years old with PhDs, working in academia at a mid-to-late career stage. Most reported working over 40 hours per week. These results are summarized in **Table 1**.

**Table 1:** Respondent Demographics

| Factor (N=874) |  | Percent | Factor (N=874) |  | Percent |
| --- | --- | --- | --- | --- | --- |
| Gender<br>(2% No Answer) | Male | 65% | Degree Type<br>(1% No Answer) | PhD | 79% |
|  | Female | 32% |  | MD | 22% |
|  |  |  |  | Other | 6% |
| Age<br>(3% No Answer) | Under 30 | 0% | Type of Work<br>(2% No Answer) | Academia | 80% |
|  | 30-39 | 2% |  | Government | 6% |
|  | 40-49 | 22% |  | Industry | 4% |
|  | 50-59 | 31% |  | Other | 9% |
|  | 60+ | 42% |  |  |  |
| Race/Ethnicity<br>(5% No Answer) | Caucasian | 76% | Career Stage<br>(3% No Answer) | Early | 3% |
|  | Asian | 10% |  | Mid | 26% |
|  | Latino | 4% |  | Late/Tenured | 59% |
|  | Black | 1% |  | Emeritus | 9% |
|  | Native American/Hawaiian | 0.1% |  |  |  |
|  | Other | 3% |  |  |  |
|  |  |  | Hours Worked per Week<br>(8% No Answer) | 40 Hours | 9% |
|  |  |  |  | 40-50 Hours | 23% |
|  |  |  |  | 50-60 Hours | 33% |
|  |  |  |  | 60-70 Hours | 18% |
|  |  |  |  | 70+ Hours | 9% |

### Peer Review Participation

The majority of respondents (76%; N=667) had served on a peer review panel in the last 3 years while 24% (N=207) had not. For those who reviewed, there was a bimodal distribution, where most respondents either served on 2-3 panels in the last 3 years or served on 7 or more panels in the last 3 years (**Figure 1a**). Federal funding agencies like the NIH have standing study sections with multiple meetings per year that typically involve multi-year appointments of their members, which may explain the bimodal participation from reviewers; the more active reviewers may have be appointed study section members (Amero, 2015). This sub-group that reviews on at least 7 panels over the 3-year timeframe will be referred to as Rev7 hereafter in this manuscript.

**Figure 1.**
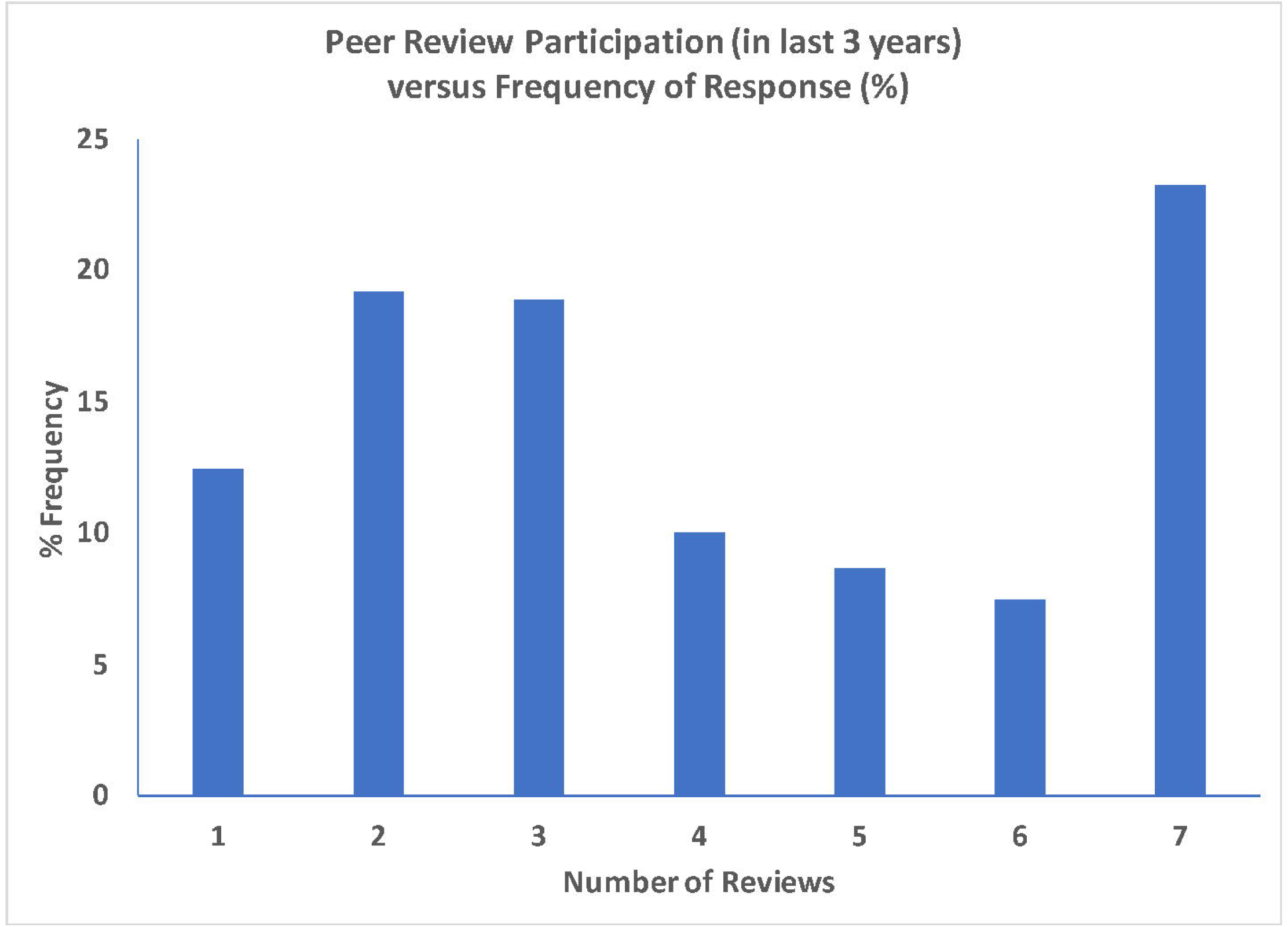
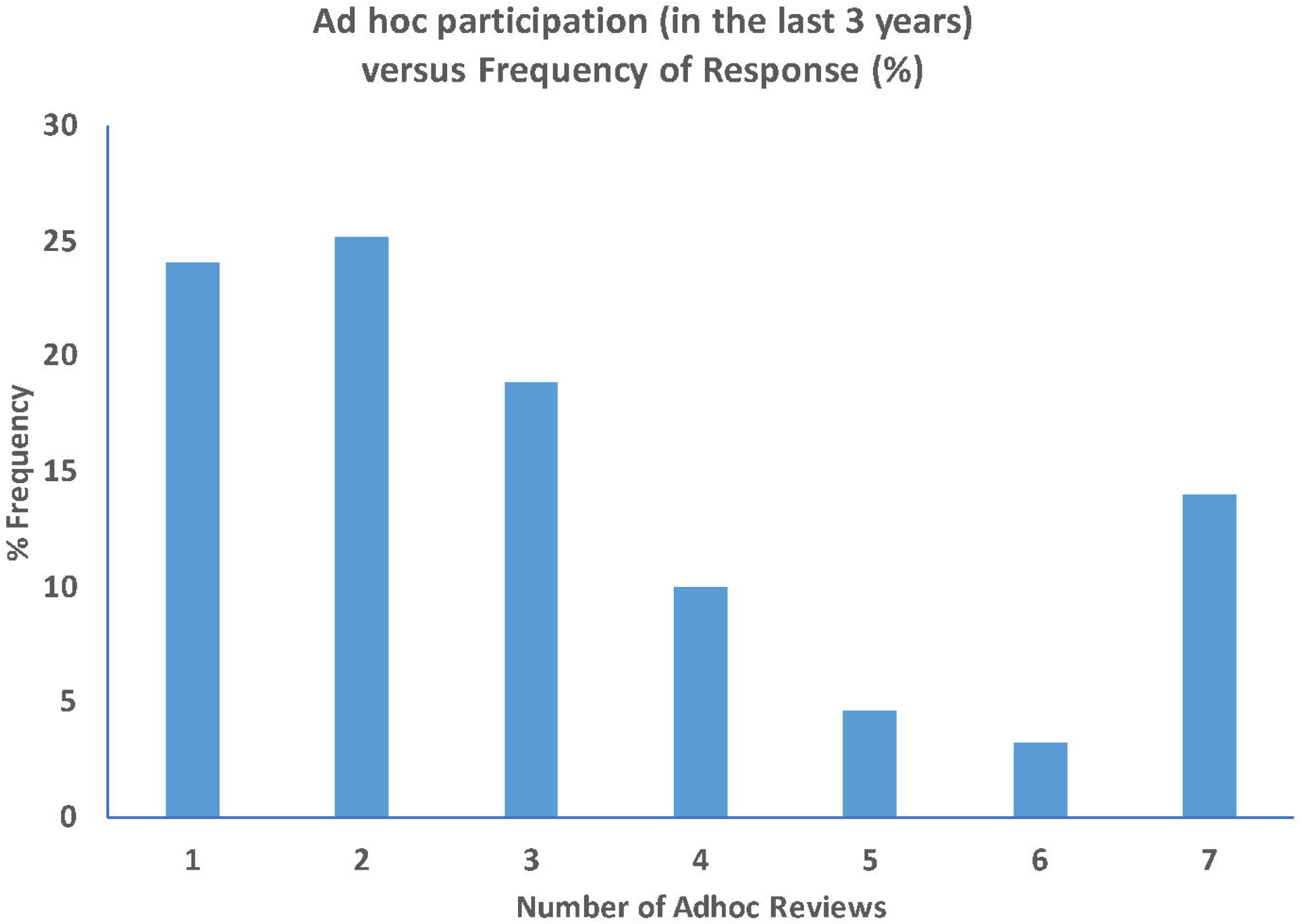
Histogram of panel and ad-hoc review participation. **a.** Level of grant peer review panel participation in the last three years (number of panels) versus frequency of response (proportion of total panel reviewer respondents; N=667) **b.** Level of ad-hoc grant peer review participation in the last three years (number of ad-hoc reviews; N=520) versus frequency of response (proportion of total ad-hoc reviewer respondents).

Of the total 2660 review panels that were reported from this survey, it was found that the Rev7 subset of reviewer respondents (23%; N=155) served on 40% of the panels (**Figure 1a**) and the remaining 77% (N=512) of reviewer respondents served on 60% of the panels, with an average review load of 3.1 ± 0.07 panels over the 3-year period. Thus, there is a significant degree of inequity observed in the distribution of respondent workload, with the top 10% reviewing 3 times the amount of the bottom 40% of respondents (Palma ratio = 3.0). This is best illustrated in the Lorenz curve in **Figure 2**. It should be noted that it is assumed that all of these reported panels are unique.

**Figure 2.**
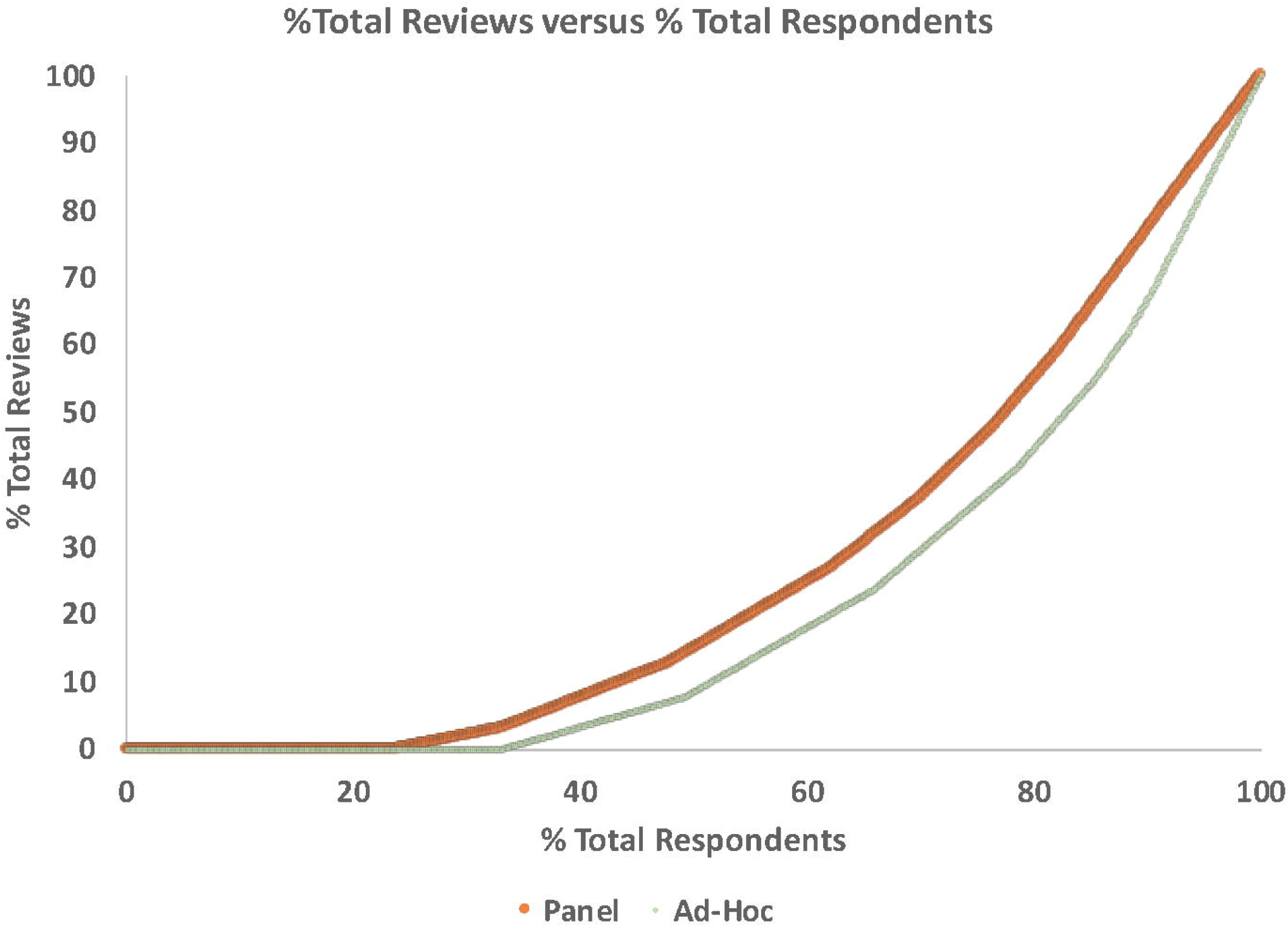
Lorenz curve for reported panel and ad-hoc reviews. The cumulative proportion of total reported reviews versus the proportion of total respondents (N=874), ordered by those who reviewed the least to those who reviewed the most. This was plotted for both panel reviews (proportion of 2660 total panel reviews in orange) and ad-hoc reviews (proportion of 1622 total ad-hoc reviews in green).

A total of 59% of respondents (N=520) indicated that they had served as an ad hoc reviewer (usually reviewing telephonically in a panel meeting setting) during the last 3 years (98 provided no answer). Again, for those who reviewed, there was a bimodal distribution, where most respondents either reviewed 2-3 times in the last 3 years or more than 6 times in the last 3 years (**Figure 1b**). The level of ad-hoc reviewing was somewhat correlated with full panel reviewing (R^2^=0.17, P<0.001). Of the 1622 ad-hoc reviews reported, 80% were performed by 38% of reviewers (4.4 ± 0.1 ad-hoc panels over the 3-year period). Again, this shows a significant degree of inequity in review workload (Palma Ratio = 9.7; **Figure 2**).

The demographic make-up of Rev7 reviewers was very similar to that of all respondents (**Table 1**), with respect to gender, age, ethnicity/race and hours worked per week. However, Rev7 respondents were more likely to have a PhD (89%), and slightly more likely to be from academia (87%). In addition, Rev7 respondents were also almost exclusively mid-to-late career stage, with no early career stage scientists and only 2.6% emeritus.

### What Predicts Grant Review Participation

In order to examine the effect of demographic variables on review panel participation, a multiple linear regression model was applied, including age, career stage, gender, race, work hours, degree, organization, as well as frequency of journal reviewing and grant submission (**Table 2**). As some respondents chose not to answer some demographic questions, any incomplete responses were removed from this analysis. Overall, the model was significant (R^2^=0.17, p<0.001, N=718); career stage (p=0.001), journal reviewing (p<0.001) and grant submission (p<0.001) were all found to be significant predictors of panel review participation. These 3 factors are examined more closely below. However, gender, age, race, degree, organization and work-week hours were not significant predictors (**Table 2**).

**Table 2.** Overall Multiple Regression Model (N=718, R^2^=0.17, p<0.001)

| Factor | Coefficient (standard error) | p-value |
| --- | --- | --- |
| Gender (Female=1)<sup>a</sup> | -0.1 (0.19) | 0.60 |
| Age (30-70 yrs)<sup>b</sup> | 0.00 (0.01) | 0.71 |
| Race (Non-Caucasian=1)<sup>a</sup> | 0.01 (0.19) | 0.95 |
| Degree (Non-PhD=1)<sup>a</sup> | -0.30 (0.23) | 0.19 |
| Organization (Non-Academic=1)<sup>a</sup> | 0.16 (0.26) | 0.53 |
| Work Week Hours (40-70 hrs)<sup>b</sup> | 0.02 (0.01) | 0.11 |
| Career Stage (Early/Mid=1)<sup>a</sup> | -0.75 (0.23) | 0.001** |
| Grant Submissions (0-7)<sup>b</sup> | 0.26 (0.04) | <0.001** |
| Journal Reviewing (0-7)<sup>b</sup> | 0.21 (0.05) | <0.001** |

Categorical variable<sup>a</sup> versus continuous numeric variable<sup>b</sup> are indicated in the table. Categorical values that equal 1 are in parentheses (e.g. Female=1).

### Grant Submission Levels

Just as with reviewing, the majority (80%; N=701) of respondents had submitted a grant in the last 3 years while only 19% (N=166) had not (7 provided no answer). Of the grant submitters, 90% indicated review participation, compared to 53% for non-submitters (X^2^(1)= 124.8; p<0.001; **Table 3**). Also, reviewer respondents submitted nearly double the number of grant applications (3.9 ± 0.1) compared to non-reviewers (2.4 ± 0.2), with Rev7 respondents submitting the most (**Table 3**). Similar results were obtained for ad-hoc reviewing, where 67% of submitters indicated ad-hoc review participation compared to 44% of non-submitting respondents (**Table 3**).

**Table 3.**

| Grant Submission | N | % Ad-hoc Reviewers | % Panel Reviewers | Number of Submission (non-Rev7) | Number of Submission (Rev7) | Rev7 vs non-Rev7 Submission Levels |
| --- | --- | --- | --- | --- | --- | --- |
| Yes | 701 | 67% (N=467) | 90% (N=628) | $3.9 \pm 0.1$ | $5.2 \pm 0.2$ | $t[665]=5.34$; $p<0.001^{**}$ |
| No | 166 | 44% (N=73) | 53% (N=88) | n.a. | n.a. | n.a. |

Of those who reported submitting grants (both reviewers and non-reviewers), 38% (N=263) indicated that their last grant submission was funded while 60% (N=418) were not (20 preferred not to answer). Similar success levels were found in the Rev7 group (40%; N=57). Review activity was not observed to be correlated with recent funding success, as 85% of those who were funded (N=263) and 82% of those who were not funded (N=418) served as reviewers.

### Reviewers of Both Manuscripts and Grant Applications

A total of 91% (N=799) of respondents had reviewed for a journal in the past 3 years (11 provided no answer) and the vast majority (75%) had reviewed 7 or more journal submissions in that time frame. It should be noted that for those who did not review for journals, there was nearly triple the proportion of Emeritus respondents (24%) as compared to the global proportion of 9% (**Table 1**; X^2^[1]= 13.7; p<0.001). Of those who reviewed for journals, 78% also participated in grant panel review, compared to 56% of those who did not review for journals (X^2^[1]= 16.4; p<0.001). Similarly, 68% of those who reviewed for journals also participated in ad-hoc review, compared to 53% of those who did not review for journals, although the difference was not highly significant (X^2^[1]= 3.849; p<0.05). When asked which is a higher personal priority (grant review or journal review), of the respondents who had indicated serving as both a grant and journal reviewer in the past 3 year, 88% viewed grant review as having equal or higher priority to journal review.

### Career Stage

Career stage can be further broken out for reviewers into the 4 groups listed in **Table 4**. Respondents who indicated participation in grant panel review had significantly lower proportions of early and emeritus career stage scientists but higher proportions of late career stage scientists compared to non-reviewers (**Table 4**). For those who reviewed, significant differences in the levels of review participation were also found across groups, with late career reviewers shouldering the largest load (F[3,653]=5.85; p<0.001; **Table 4**). No early career stage scientists at all were included in Rev7, and only 2.6% emeritus (n=4) were, while late career stage scientists increased in proportion to 73% (n=112) and mid-level was stable at 24% (n=37). It is noted that only 37% of emeritus reviewers reported submitting any grants (compared to 80% for the whole sample), at a rate of 0.99 ± 0.2 grants in the 3-year period.

**Table 4.**
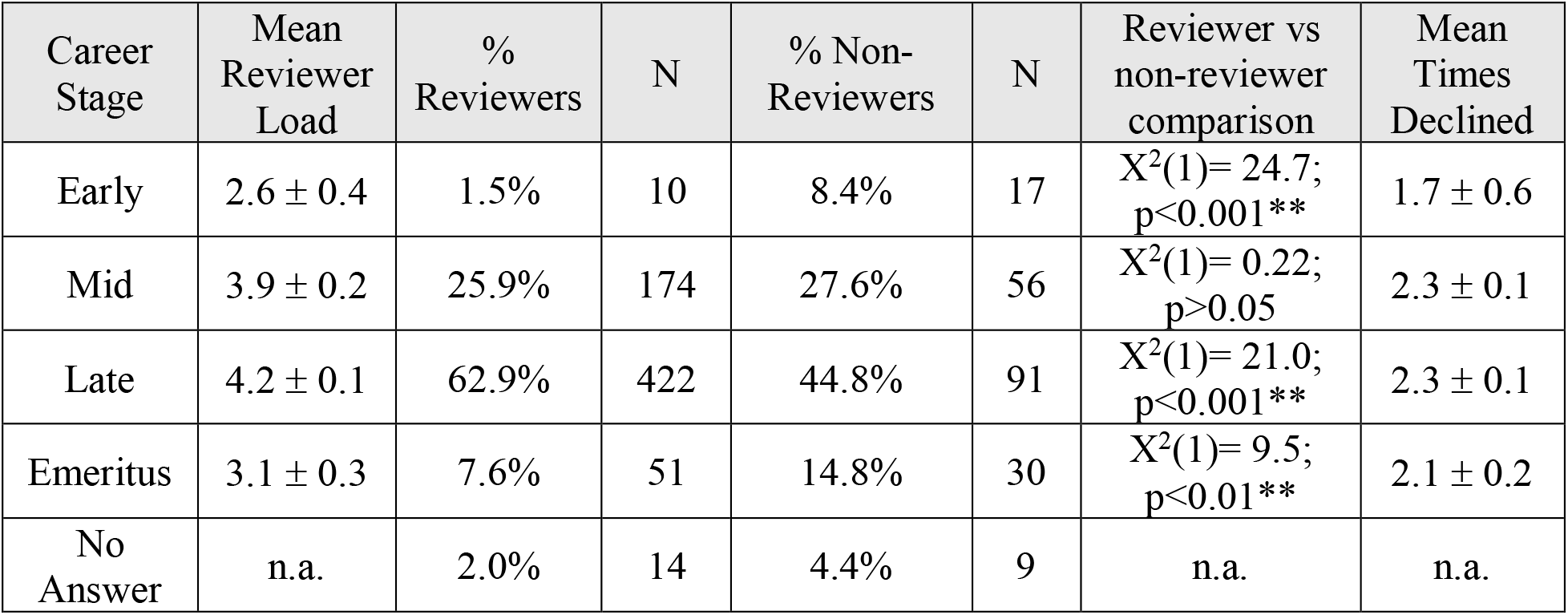

Respondents who indicated participation in ad-hoc review were comprised of significantly lower proportions of early and emeritus career stage scientists but higher proportions of late career stage scientists compared to non-reviewers (data not shown). The levels of participation for ad-hoc reviewers showed a similar relationship to career stage (F[3,509]=4.76; p=0.003), where late career reviewers participated in more ad-hoc reviews than the other reviewer groups (**Table 4**).

### Reasons to Participate in Peer Review

When reviewer respondents were asked to select the reasons (choose all that apply) for accepting an invitation to serve on a peer review panel, 82-90% of respondents selected giving back to the scientific community, which was overwhelmingly the most popular motivation (**Table 5**). Interestingly, 51% of non-Rev7 respondents selected informing their own grantsmanship compared to 67% of Rev7 respondents, which is consistent with the elevated levels of grant submission among Rev7 respondents (**Table 5**). Surprisingly, only 18% selected expectation from the funding agency, for both Rev7 and non-Rev7 reviewers.

**Table 5.**

| Motivation to Review or Impact on Career | Non-Rev7 (%/N) | Rev7 (%/N) | Chi Squared |
| --- | --- | --- | --- |
| Giving back to the scientific community | 82% / 420 | 90% / 139 | $X^2[1] = 5.7; p < 0.05^*$ |
| Gaining exposure to new and innovative scientific areas | 54% / 278 | 60% / 93 | $X^2[1] = 1.7; p > 0.05$ |
| Informing your own grantsmanship | 51% / 263 | 67% / 104 | $X^2[1] = 12.4; p < 0.001^{**}$ |
| Networking opportunities | 37% / 188 | 44% / 68 | $X^2[1] = 2.7; p > 0.05$ |
| Enhancing your career/resume | 26% / 132 | 30% / 46 | $X^2[1] = 1.0; p > 0.05$ |
| Expectation from the funding agency | 18% / 94 | 18% / 28 | $X^2[1] = 0.003; p > 0.05$ |
| Honorarium | 9% / 44 | 9% / 14 | $X^2[1] = 0.04; p > 0.05$ |
| Serving as a reviewer on peer review panels had positively impacted career | 87% / 434 | 92% / 142 | $X^2[1] = 3.1; p > 0.05$ |
| Influenced career through improvements in grantsmanship | 66% / 341 | 75% / 116 | $X^2[1] = 4.1; p < 0.05^*$ |
| Influenced career through increased exposure to new scientific ideas | 61% / 316 | 68% / 106 | $X^2[1] = 1.6; p > 0.05$ |
| Influenced career through improved networking/collaboration opportunities | 40% / 204 | 56% / 87 | $X^2[1] = 13.2; p < 0.001^{**}$ |

Although only 27% of all reviewer respondents were motivated to enhance their career, 87-92% felt that serving as a reviewer on peer review panels had positively impacted their career, most frequently through improvements in grantsmanship. Differences were observed between Rev7 and other reviewers regarding perceptions of networking/collaboration opportunities provided by peer review panels, with the Rev7 group viewing it more positively.

### Declined Peer Review Participation

When asked how many times reviewers had declined an invitation to serve on a peer review panel in the past 3 years, respondents indicated they declined on average 2.3 ± 0.1 times (N=618) during this time period. The declination rate was the same for reviewing and non-reviewing respondents; 2.2 ± 0.2 times (N=78) for non-reviewers versus 2.2 ± 0.1 times (N=396) for non-Rev7 reviewers, although there was a slight increase for Rev7 respondents; 2.6 ± 0.1 times, N=139. Career stage did not seem to be a factor either (**Table 4**; F[3,602]=0.97; p=0.40). Small but significant differences in declination (t[613]=3.6; p<0.001) were found between grant submitters (2.4 ± 0.1 times) and non-submitters (1.7 ± 0.1 times). It is possible funding agencies may be more inclined to invite former applicants to review; however no differences were found between recently successful and unsuccessful submitters (t[516]=1.9; p=0.06).

Of those who indicated they declined, 60% (N=368) indicated this was due to limited time, 28% (N=172) due to personal reasons (like holiday, sickness, travel), and 21% (N=129) due to the review timeline being too compressed. Only 21% (N=129) indicated declining was due to poorly matching expertise, 12% (N=75) due to conflict of interest, and 1% (N=11) due to issues with the funding agency (respondents could select all that apply). There were no significant differences in reasons for declination between non-reviewers and Rev7 and non-Rev7 reviewers.

### Estimated Maximum Review Capacity

Significant differences were seen across non-reviewers, reviewers, and Rev7 respondents in terms of the maximum number of peer review panels they would be willing to participate in per year (F[2,795]=147; p<0.001). Non-Rev7 reviewers indicated a willingness to serve on no more than 2.0 ± 0.04 panels per year (N=502, 14 provided no answer), and Rev7 reviewers indicated no more than 3.1 ± 0.06 panels per year (N=150, 5 provided no answer). The maximum preferred review loads for both groups were larger than their actual levels of participation (**Supplementary Table 1**).

Using the data in **Supplementary Table 1** in **Appendix A**, one can get a sense of annual hourly commitment to panel reviews and how this actual commitment relates to maximum capacities. It should be noted that these figures are based on the self-reported totals of 55 and 57 hours per week, for non-Rev7 and Rev7, respectively. It is estimated that non-Rev7 reviewers spent an average of 47 ± 2 hours per year in grant review (1.7% of total annual hours), and had a maximum preference of 83 ± 3 hours per year (3.1% of total annual hours). For Rev7 reviewers, however, it is estimated they spent nearly three times as many hours, with an average of 132 ± 3 hours per year in grant review (4.8% of total annual hours), and had a nearly double maximum preference of 151 ± 5 hours per year (5.4% of total annual hours). If one takes the maximum preferred values as a threshold of possible reviewer involvement, it is calculated that non-Rev7 reviewers are reviewing at 56% of their estimated threshold while Rev7 reviewers are reviewing at 87% capacity. In terms of the total 36,751 hours actually spent on the 2199 panels included in this analysis, the 145 Rev7 reviewers spent a total of 18,279 hours participating in 973 panel meetings, while the 390 non-Rev7 reviewers spent almost the same amount of total hours (18,472 hours) in 1226 panel meetings.

## Discussion

### Survey Generalizability

While the overall response rate for this analysis and the previous published analysis was the same (9.4% and 1231 responses), the number of respondents included in this analysis is different than previously published (Gallo 2018). This is because in our previous publication, only 850 had complete responses to questions relevant to that analysis, whereas in this analysis, 874 had complete responses to questions 2efg, and were therefore included. While our response rate was low, it is similar to surveys of journal peer review, which had rates of 8-10% (Ware and Monkman, 2008; Sense About Science, 2009).

Most of our 874 survey respondents were white males with PhDs who were 40 years or older, working in academia, at their mid-to-late career stage, and working more than 40 hours per week (**Table 1**). These demographics are similar to those reported for NIH reviewers (NIH, 2012, Rockey, 2015) in terms of gender, race, and degree. However, female scientists are underrepresented in our respondent population (32%) compared to all life scientists (48%) as are scientists of Asian descent (10%) compared to all biochemists, biophysicists, and molecular biologists (22%), while white men are overrepresented (National Science Foundation [NSF] 2015; DataUSA, 2016). Also, our respondents tended to be older than NIH reviewers, with an overrepresentation of respondents over 60 in our survey (42%) compared to NIH reviewers (16%; NIH, 2012). That said, the career stages of our respondents were very similar to those of NIH study section panelists (NIH 2008).

The overwhelming majority of our survey respondents had recently submitted grants, with a 38% reported success rate of funding (similarly 40% for Rev7). As these rates are higher than the overall funding success rates reported by the National Science Foundation (24%; NSF 2018) and NIH (19%; Lauer 2018), it is likely our survey population has a reasonable amount of funding, which is in line with the funding status of NIH peer reviewers (Rockey, 2015). Thus, it is felt that this sample is roughly similar to the make-up of most NIH research panels with respect to funding. However, it should also be mentioned that review activity was not observed to be directly associated with recent funding success.

### Participation Levels in Grant Peer Review

The majority of our survey respondents had served on a grant peer review panel during the past 3 years, although a substantial minority had not (24% for panel reviews and 40% for ad-hoc reviews). There was a bimodal distribution in the level of reviewer participation observed for both panel and ad-hoc reviewing (**Figure 1**). The Rev7 respondents, identified as the high end of the participation distribution, are less likely to be emeritus or early-stage career scientists and more likely to submit more grant applications than non-reviewers or non-Rev7 reviewers. Given their workload, many of these Rev7 respondents may have been appointed to a federal study section (Amero, 2015). Unfortunately, information was not collected related to the level of current funding and from what agency, which would have allowed further definition of this population.

Nevertheless, substantial inequities in review participation were quantified in the Palma ratios of 3.0 and 9.7 for panel and ad-hoc reviews, respectively. It is probable that respondents reviewed for agencies from which they have received funding and it is likely that the distribution of review participation observed here is due to the variety of funding agencies to which the respondents had submitted. Many smaller agencies have much less frequent peer review meetings and may be less likely to convene standing panels, which may reduce the number of times these respondents are invited to participate. For instance, for Department of Defense research programs, proposal receipt and peer review usually occur on an annual cycle (CDMRP 2018). Again, funding agency information was not collected from respondents (e.g. where they submitted grants and where they participated in reviews).

### Review Capacity

Based on respondents’ indications of their preferred maximum panel review load, it was calculated that Rev7 respondents are already working at 87% of their maximum preferred hours dedicated toward grant review (compared to non-Rev7 reviewers at 56%). In addition, it was found that Rev7 respondents submitted 33% more grants during this time-period than non-Rev7 respondents. Thus, it is likely that Rev7 respondents are particularly burdened by the grant submission/grant review cycle. If Rev7 reviewers are federal study section members, they may complete their appointment in a few years; nevertheless, what effect this time-intensive commitment may have on the sustainability of the grant peer review system is unclear. Considerable reviewer time fatigue in this population is likely (which may affect review quality), and if positive funding status is a requirement for participation, the size of the reservoir of eligible reviewers who can replace the Rev7 reviewers is unclear, especially given the concentration of funding among a relatively small group of researchers (Wahls, 2018). This result highlights the need to understand the motivations and preferences of potential reviewers.

### Predictors and Motivators of Review Participation

Multiple regression analysis suggests gender, age, race, degree, organization and hours worked per week did not significantly predict reviewer participation, although career stage, journal reviewing, and grant submission did. The correlation of review participation with career stage was likely driven by the low levels of early stage and emeritus reviewer participation. The lower levels of participation in early stage reviewers were likely due to the fact that they were not invited, due to their short track record (as suggested by NIH panel demographics; NIH 2012, Rockey, 2015) whereas emeritus reviewers may not review in part because they submit fewer, if any, grants (only 37% reported submitting compared to 80% for the whole sample. Similarly, the lack of journal reviewing was probably correlated to the lack of panel reviewing due to the high concentration of emeritus respondents.

The level of grant submission was also a significant predictor of review activity, which coincides with the most popular review motivation, giving back to the scientific community. This motivation is likely to be enhanced when a scientist is invested in the research funding cycle and others are reviewing their grants. Interestingly, individual recent funding success was not associated with review participation. It is likely the sense of duty was due to having their own grant submissions reviewed, and not as direct reciprocity to the funding agency for receiving research awards. While respondents may be eligible and expected to be study section panelists due to their funding, only 18% indicated expectation from the funding agency as a motivator to review, which was true for both Non-Rev7 and Rev7 populations. Together, these findings are consistent with the most popular responses reported in surveys on journal review (Ware 2008; Nobarany, 2015; SAS, 2009; and Ware and Monkman, 2008), as well as surveys of European grant review (Schroter, 2010) and may represent a common sense of duty to the scientific gatekeeping function of peer review, and the preferred way to fulfill expectations to service the profession among those in academia. The desire to sustain the system they are utilizing as grant applicants most likely explains why the vast majority of our respondents viewed grant review as equal or greater priority to journal review.

The next two most popular reasons our respondents selected for accepting an invitation to serve on a peer review panel were exposure to new and innovative scientific areas and informing grantsmanship, both of which were reported at a higher rate in the Rev7 sample compared to other reviewer respondents. These were also the two most popular reasons selected by our respondents for how serving as a reviewer on peer review panels had positively impacted their career. These are likely to be important incentives for potential reviewers in their early career stages. In fact, one of the outlined benefits listed in the NIH CSR Early Career Reviewer program is to “improve grant writing skills by getting an insider’s view of how grant applications are evaluated (NIH 2018).”

Interestingly, differences were observed between Rev7 and other reviewers regarding perceptions of networking/collaboration opportunities provided by peer review panels, with the Rev7 group viewing it more positively. This is perhaps because study sections meet more often and have relatively consistent rosters, which provides a greater opportunity to network among the same group than ad-hoc style review panels

Overall, review participation did not correlate well with declining to review, and across all respondent groups, and the reasons for declining to participate were largely the same—a lack of time. Interestingly, this result is similar to a recent survey of European grant reviewers, who indicated factors relating to time constraints were the biggest barriers to undertaking grant review (Schroter, 2010). Given the time intense nature of grant writing (some have estimated an average of 34 days [272 hours] to write each application; Herbert, 2013), future studies should examine closely the time constraints of scientists and their relationship to the sustainability of the peer review system; one study found 42% of sampled scientists feel they have sacrificed work-life balance for a career in science (Woolsten, 2016).

### Conclusions

In conclusion, the results of this independent survey suggest participation in grant peer review is uneven, with some groups much closer to their maximum capacity than others. Most are motivated by giving back to a funding system with which they are engaged as an applicant; the less engaged they are with the system, the less compelled they likely feel to provide review services for it. New incentives and motivations will likely be important to enlarge the pool of reviewers, not only in terms of the sustainability of the system, but also to increase recognition and engagement in an important gatekeeping process in science, and to add diversity to improve overall review quality.

## Appendix A

### Reviewer Preferences versus Experience

When asked the actual number of days of their last peer review panel meeting, non-Rev7 reviewers indicated 1.8 ± 0.03 days, which compared well to their preference of 2.0 ± 0.03 days, and similar results were found for Rev7 reviewers (1.9 ± 0.05 actual days versus the preference of 2.1 ± 0.05 days). When asked the actual number of R01-type grant applications they were assigned at the last peer review panel meeting, non-Rev7 reviewers reported a smaller review load (5.6 ± 0.1) than Rev-6 reviewers (6.9 ± 0.2; t[570]=6.1; p<0.001), although these values are similar the average 7 assignments that have been reported in NIH study Sections (NIH 2008). Non-Rev7 reviewers indicated a clear preference for fewer assignments, 4.5 ± 0.08 compared to actual load (t[848]=8.0; p<0.001), as did Rev7 reviewers, 5.4 ± 0.1 compared to actual load (t[292]=7.4; p<0.001). The amount of time spent per assignment reviewing each application before the meeting was fairly similar between non-Rev7 (4.5 ± 0.08 hours per application) and Rev7 reviewers (5.0 ± 0.08 hours per application), with only small differences detected (t[569]=3.2; p=0.002).

When asked about their preferred mode of peer review (face-to-face, video/audio teleconference, internet-assisted or other), 77% (N=382) of non-Rev7 reviewers indicated face-to-face, despite that only 44% (N=221) of recent reviews being face-to-face (X^2^[1]=110, p<0.001). Similarly, 89% (N=133) of Rev7 reviewers preferred face-to-face panel meetings compared to 68% (N=102) of recent reviews using this mode of review (X^2^[1]=18.9, p<0.001). When asked to rate on a Likert scale of 1-5 (1 most influential, 5 least influential) the reasons that influence their selection of preferred panel meeting format, the average rating for non-Rev7 and Rev7, respectively, was 2.2 ± 0.06 and 1.9 ± 0.1 for level of communication among panel members, 2.8 ± 0.06 and 2.6 ± 0.1 for networking opportunities, 2.8 ± 0.05 and 2.8 ± 0.1 for likelihood to participate, and 3.0 ± 0.06 and 3.0 ± 0.1 for logistical convenience. It should be noted that for reviewers who prefer teleconferencing panels, they feel more strongly about logistical convenience (2.4 ± 0.2) then they do about the level of panel communication (3.0 ± 0.06).

**Supplementary Table 1.**
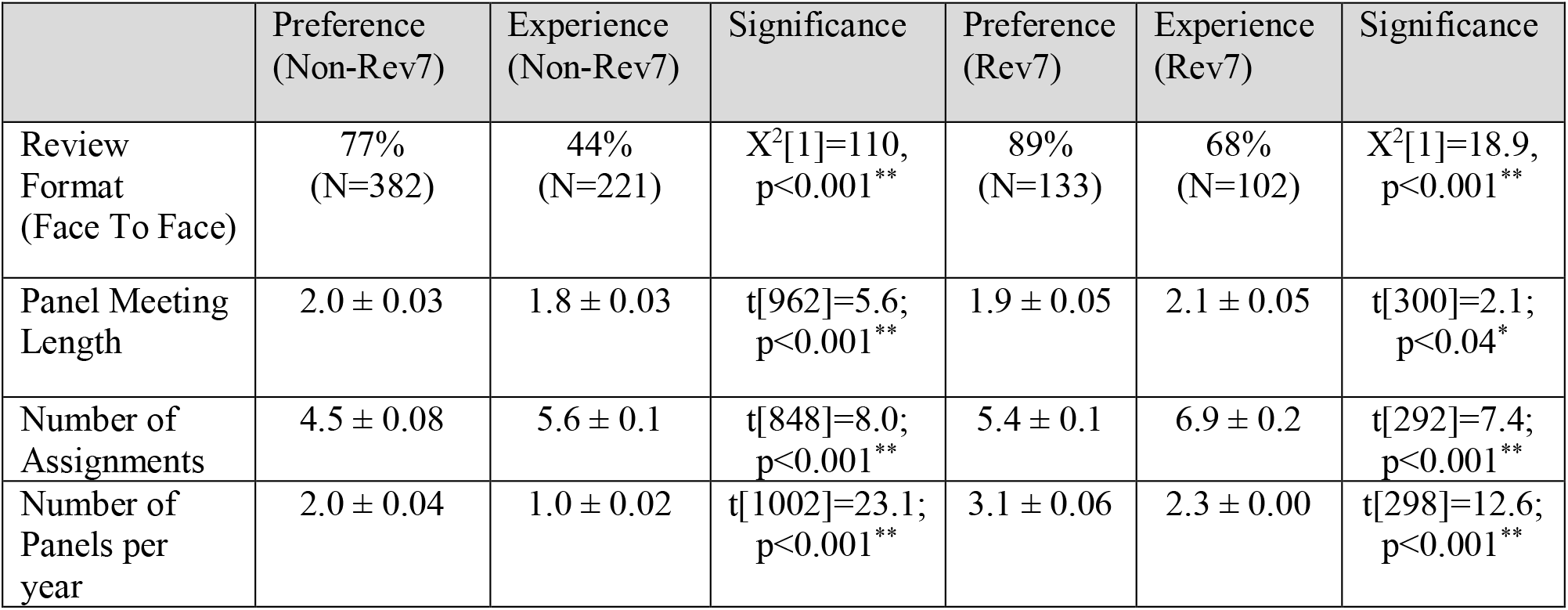

|  | Preference<br>(Non-Rev7) | Experience<br>(Non-Rev7) | Significance | Preference<br>(Rev7) | Experience<br>(Rev7) | Significance |
| --- | --- | --- | --- | --- | --- | --- |
| Review<br>Format<br>(Face To Face) | 77%<br>(N=382) | 44%<br>(N=221) | $X^2[1]=110$,<br>$p<0.001^{**}$ | 89%<br>(N=133) | 68%<br>(N=102) | $X^2[1]=18.9$,<br>$p<0.001^{**}$ |
| Panel Meeting<br>Length | $2.0 \pm 0.03$ | $1.8 \pm 0.03$ | $t[962]=5.6$;<br>$p<0.001^{**}$ | $1.9 \pm 0.05$ | $2.1 \pm 0.05$ | $t[300]=2.1$;<br>$p<0.04^*$ |
| Number of<br>Assignments | $4.5 \pm 0.08$ | $5.6 \pm 0.1$ | $t[848]=8.0$;<br>$p<0.001^{**}$ | $5.4 \pm 0.1$ | $6.9 \pm 0.2$ | $t[292]=7.4$;<br>$p<0.001^{**}$ |
| Number of<br>Panels per<br>year | $2.0 \pm 0.04$ | $1.0 \pm 0.02$ | $t[1002]=23.1$;<br>$p<0.001^{**}$ | $3.1 \pm 0.06$ | $2.3 \pm 0.00$ | $t[298]=12.6$;<br>$p<0.001^{**}$ |

## Appendix B

### Full Survey

#### Section 1: Demographics

[]What is your gender? Please choose all that apply:

Male

Female

Prefer not to answer

[]What is your age? Please choose only one of the following:

Under 30

30–39

40–49

50–59

60+

[]Please specify your race/ethnicity

Please choose all that apply:

American Indian or Alaska Native

Asian or Asian American

Black or African American

Hawaiian or Other Pacific Islander

Hispanic or Latino

Non-Hispanic White/Caucasian

Other

Prefer not to answer

[]What type of degree(s) do you have?

Please choose all that apply:

PhD or other research doctorate

MD

DDS

DVM or VMD

Other

Prefer not to answer

[]What type of an organization do you work for?

Please choose only one of the following:

Academia

Government

Industry

Other

[]What stage of career have you reached?

Please choose only one of the following:

Early career

Mid career

Late career/tenured

Emeritus

[]On average, how many hours do you work each week?

Please choose only one of the following:

40 h

40–50 h

50–60 h

60–70 h

70 + h

[]Please provide any comments that justify your responses under Section 1, Demographics.

#### Section 2: Grant submission and peer review experience

[]Have you submitted a grant for peer review in the last 3 years? Please choose only one of the following:

Yes

No

[]If you answered yes to submitting a grant for peer review in the past 3 years, how many grant applications have you submitted in that time frame? Please choose only one of the following:

1

2

3

4

5

6

7 or more

[]Did you receive reviewer feedback on your last grant submission? Please choose only one of the following:

Yes

No

[]Was your last application successful, i.e., were you funded? Please choose only one of the following:

Yes

No

[]Have you served on a peer review panel in the last 3 years? Please choose only one of the following:

Yes

No

[]If you answered yes to serving on a peer review panel in the last 3 years, how many peer review panels have you served on in that time frame? Please choose only one of the following:

1

2

3

4

5

6

7 or more

[]If you answered yes to serving on a peer review panel in the past 3 years, please select the mode of your last peer review panel meeting. Please choose only one of the following:

Face-to-face

Remote (video/teleconference)

Internet-assisted

Other

[]How many ad-hoc reviews (usually one or two grant applications reviewed telephonically that are being evaluated in a panel meeting setting) have you performed in the past 3 years? Please choose only one of the following:

0

1

2

3

4

5

6

7 or more

[]Have you reviewed for a journal in the last 3 years?

Please choose only one of the following:

Yes

No

[]If you answered yes to reviewing for a journal in the past 3 years, how many submissions have you reviewed in that time frame? Please choose only one of the following:

1

2

3

4

5

6

7 or more

[]What is a higher personal priority: grant review or journal review? Please choose only one of the following:

Grant review

Journal review

Both are equal priority

Neither is a priority

[]Please elaborate on your responses under Section 2, Grant submission and peer review experience.

#### Section 3: Investigator attitudes toward grant review

If you answered yes to receiving feedback on your last grant submission, please answer Section 3 of the questionnaire.

If you answered no, please proceed to Section 4.

[]On a scale of 1–5 (1 most useful, 5 least useful), overall how useful was the reviewer feedback you received on your last grant submission? Please choose only one of the following:

1

2

3

4

5

[]On a scale of 1–5 (1 most useful, 5 least useful), how useful was the reviewer feedback in improving your grantsmanship?Please choose only one of the following:

1

2

3

4

5

[]If you were not funded, on a scale of 1–5 (1 most useful, 5 least useful), how useful was the reviewer feedback in improving your future submissions? Please choose only one of the following:

1

2

3

4

5

[]On a scale of 1–5 (1 most useful, 5 least useful), how useful was the reviewer feedback in informing your future scientific endeavors in the proposed research area?

Please choose only one of the following:

1

2

3

4

5

[]Did you feel the reviewer feedback was well written, cohesive, and balanced? Please choose only one of the following:

Yes

No

[]Did you feel the reviewer feedback was fair and unbiased? Please choose only one of the following:

Yes

No

[]Overall, in what area(s) did the reviewer feedback primarily focus? Please choose all that apply:

Potential impact of research

Hypothesis

Research methodology

Innovation potential

Preliminary data

Responsiveness to funding mechanism

Statistical issues

Qualifications of research team

Budget

Other

[]Did the reviewers comment on the riskiness of the research project?

Please choose only one of the following:

Yes

No

[]Based on the reviewer feedback you received, do you feel that the reviewers had the appropriate expertise to evaluate your grant application?

Please choose only one of the following:

Yes

No

[]Please elaborate on your responses under Section 3, Investigator Attitudes Towards

Grant review.

#### Section 4: Reviewer attitudes towards grant review

[]What are your reasons for accepting an invitation to serve on a peer review panel? Please choose all that apply:

Desire to give back to the scientific community

Networking opportunities

Informing your own grantsmanship

Gaining exposure to new and innovative scientific areas

Enhancing your career/resume

Expectation from the funding agency

Honorarium

Other

[]Do you feel that serving as a reviewer on peer review panels has positively impacted your career?

Please choose only one of the following:

Yes

No

[]If you feel that serving as a peer reviewer has positively impacted your career, in what ways has serving as a reviewer influenced your career?

Please choose all that apply:

Bolstered your career

Improved your grantsmanship

Increased your exposure to new scientific ideas

Improved your networking/collaboration opportunities

Other

[]In general, which type of panel meeting format do you prefer? Please choose only one of the following:

Face-to-face

Virtual [teleconference/videoconference]

Internet-assisted

[]On a scale of 1–5, (1 most influential, 5 least influential), please rate the following factors in influencing your selection of preferred panel meeting format:

Logistical convenience

Level of communication among panel members

Networking opportunities

Likelihood to participate on panel

[]In the last 3 years, how many times have you declined an invitation to serve on a peer review panel?

Please choose only one of the following:

1

2

3

4

5

6

7 or more

[]What were your reasons for declining an invitation to serve on a peer review panel? Please choose all that apply:

Limited free time

Poor expertise match

Personal reasons (holiday, sickness, travel)

Review timeline too compressed

Conflict of interest

Issue with funding agency

Other

[]What is the maximum number of peer review panels/committees you prefer to serve on per year?

Please choose only one of the following:

1

2

3

More than 3

[]What is the maximum number of days you prefer to attend a peer review panel meeting?

Please choose only one of the following:

1

2

3

More than 3

[]What is the maximum number of R01-type grant applications you prefer to be assigned for a peer review panel meeting?

Please choose only one of the following:

1–2

3–4

5–6

7

More than 7

[]What was the actual number of days of your last peer review panel meeting? Please choose only one of the following:

1

2

3

More than 3

[]What was the actual number of R01-type grant applications you were assigned to review at your last peer review panel meeting? Please choose only one of the following:

1–2

3–4

5–6

7–8

More than 8

[]On average, how many hours did you spend reviewing each grant application before the panel meeting?

Please choose only one of the following:

1–2

2–3

3–4

4–5

5–6

7 or more

[]Please elaborate on your responses under Sect. 4, Reviewer attitudes towards grant review.

#### Section 5: Peer review panel meeting proceedings

[]Please answer the following questions in relation to your last peer review meeting. On a scale of 1–5 (1 most definitely, 5 not at all), was your scientific expertise necessary and appropriately used in the review process?

Please choose only one of the following:

1

2

3

4

5

[]On a scale of 1–5 (1 most definitely, 5 not at all), from your perspective was the expertise of the other panel members necessary and appropriately used in the review process?

Please choose only one of the following:

1

2

3

4

5

[]Did the grant application discussions facilitate reviewer participation? Please choose only one of the following:

Yes

No

[]Were the grant application discussions fair and balanced?Please choose only one of the following:

Yes

No

[]On a scale of 1–5 (1 most useful, 5 least useful), how useful were the grant application discussions in clarifying differing reviewer opinions?

Please choose only one of the following:

1

2

3

4

5

[]On a scale of 1–5(1 extremely effective, 5 no effect), did the grant application discussions affect the outcome? Please choose only one of the following:

1

2

3

4

5

[]On a scale of 1–5 (1 most appropriate, 5 least appropriate), were the evaluation criteria appropriate to judge the best science and move the field forward?

Please choose only one of the following:

1

2

3

4

5

[]On a scale of 1–5 (1 extremely important, 5 of no importance), how important is the PI’s track record to assessing an investigator initiated (R01)-type application?

Please choose only one of the following:

1

2

3

4

5

[]In general, does a PI’s track record temper your assessment of any detected methodological weaknesses?

Please choose only one of the following:

Yes

No

[]On a scale of 1–5 (1 most definitely, 5 not at all), did the grant application discussions promote the best science? Please choose only one of the following:

1

2

3

4

5

[]Was innovation factored into selecting the best science? Please choose only one of the following:

Yes

No

[]Did you view innovation as an essential component of scientific excellence when evaluating the grant applications? Please choose only one of the following:

Yes

No

[]Did the risks associated with innovative research impact the scores you assigned to the grant applications?

Please choose only one of the following:

Yes

No

[]On a scale of 1–5 (1 completely, 5 not at all), how much did the seniority of your fellow panel members influence your evaluations during the panel deliberations?

Please choose only one of the following:

1

2

3

4

5

[]Was the format and duration of the grant application discussions sufficient to allow the non-assigned reviewers to cast well informed merit scores?

Please choose only one of the following:

Yes

No

[]On a scale of 1–5 (1 extremely useful, 5 not useful at all), how useful was the Chair in facilitating the application discussions? Please choose only one of the following:

1

2

3

4

5

[]Please elaborate on your responses under Section 5, Peer review panel meeting proceedings.

Thank you for taking the time to fill out the survey. Have a wonderful day! Thank you for completing this survey.

